# Programmable chronogenetic gene circuits for self-regulated circadian delivery of biologic drugs

**DOI:** 10.1101/2025.03.14.643274

**Authors:** Amanda Cimino, Fiona Pat, Omolabake Oyebamiji, Lara Pferdehirt, Christine T.N. Pham, Erik D. Herzog, Farshid Guilak

## Abstract

Cells of the body rely on the circadian clock to orchestrate daily changes in physiology that impact both homeostatic and pathological conditions, such as the inflammatory autoimmune disease rheumatoid arthritis (RA). In RA, high levels of proinflammatory cytokines peak early in the morning hours, reflected by daily changes in joint stiffness. Chronotherapy (or circadian medicine) seeks to delivery drugs at optimal times to maximize their efficacy. However, chronotherapy remains a largely unexplored approach for disease modifying, antirheumatic treatment, particularly for cell-based therapies. In this study, we developed autonomous chronogenetic gene circuits that produce the biologic drug interleukin-1 receptor antagonist (IL-1Ra) with desired phase and amplitude. We compared expression of IL-1Ra from circuits that contained different circadian promoter elements (E’-boxes, D-boxes, or RREs) and their ability to respond to inflammatory challenges in murine pre-differentiated induced pluripotent stem cells (PDiPSC) or engineered cartilage pellets. We confirmed that each circuit reliably peaked at a distinct circadian time over multiple days. Engineered cells generated significant amounts of IL-1Ra on a circadian basis, which protected them from circadian dysregulation and inflammatory damage. These programmable chronogenetic circuits have the potential to align with an individual’s circadian rhythm for optimized, self-regulated daily drug delivery.

## Introduction

The circadian clock is a fundamental cellular mechanism that orchestrates biological oscillations with approximately 24-hour periods. The suprachiasmatic nucleus acts as a central clock in the vertebrate brain that synchronizes to environmental cues (e.g., light and dark) to drive daily rhythms in physiology and behavior [1]. Peripheral circadian clocks in many tissues entrain to daily signals such as endocrine, neural, and mechanical cues [2, 3]. At a cellular level, autoregulated transcriptional and translational feedback loops generate circadian rhythms in gene expression. The core circadian feedback loop is controlled by dimers of BMAL1 (basic helix-loop-helix ARNT-like protein 1) and CLOCK that bind short promoter regions called E-boxes upstream of clock genes to activate their expression, which includes the PER (period) and CRY (cryptochrome) proteins [2]. PER:CRY proteins dimerize and interact with BMAL1:CLOCK to repress their transcriptional activity, thereby regulating their own expression. These proteins are eventually degraded, and the cycle continues with an approximately 24-hour period. Additional feedback loops driven by protein binding to D-boxes and Rev-Erb/ROR response elements (RREs) coordinate the precise timing of circadian gene and protein expression and further stabilize the cellular-level circadian clock [4] (**Fig. 1**).

**Figure 1.**
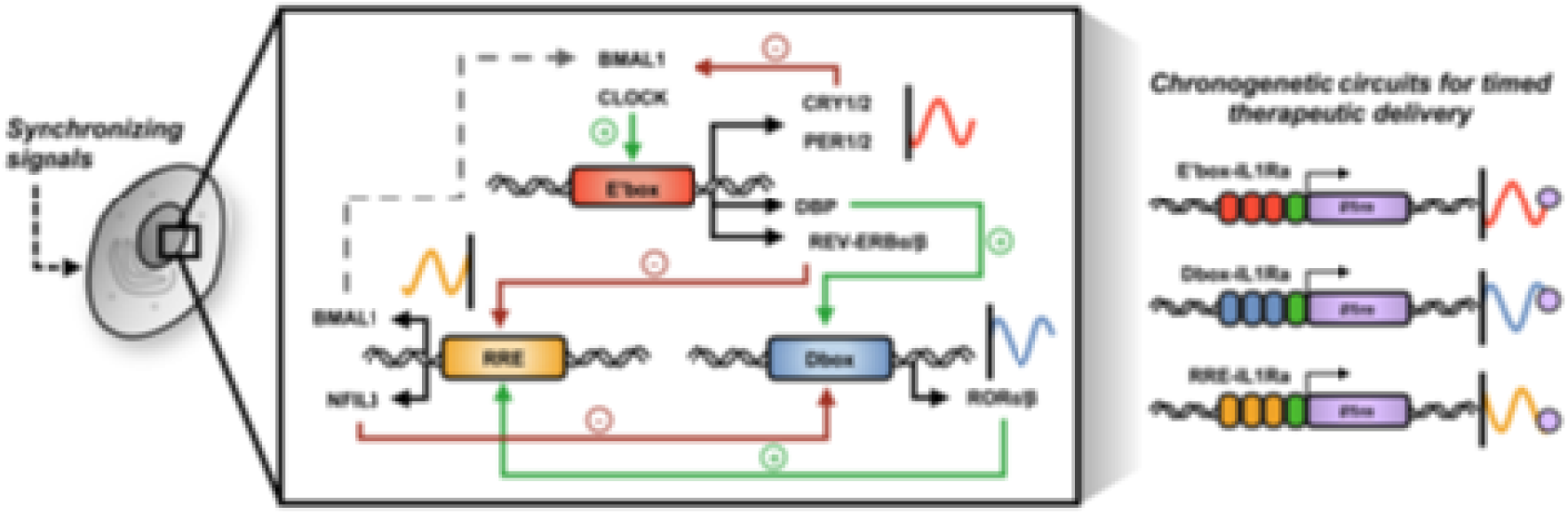
Overview of synthetic chronogenetic gene circuits. External cues synchronize circadian rhythms throughout the body, and at a cellular level, interacting feedback loops genetically regulated by clock-controlled elements (i.e., E’-boxes, D-boxes, and RREs) coordinate internal cellular timing based on their unique expression phases across a 24-hour period. By harnessing this feature of their expression, we generated synthetic chronogenetic gene circuits that cyclically generate the therapeutic IL-1Ra at different phases throughout a day for programmable therapeutic delivery. BMAL1: Basic Helix-Loop-Helix ARNT Like 1; CLOCK: Clock Circadian Regulator; CRY: Cryptochrome Circadian Regulator; PER: Period Circadian Regulator; DBP: D-Box Binding PAR BZIP Transcription Factor; REV-ERB: Nuclear Receptor Subfamily 1 Group D Member 1; ROR: Retinoic Acid Receptor-Related Orphan Receptor; NFIL3: Nuclear Factor, Interleukin 3 Regulated.

In addition to their role in normal physiologic processes in the body, it is now clear that circadian rhythms also play critical roles in pathologic conditions such as sleep disorders, cancer, metabolic diseases, neurodegenerative diseases, and autoimmune diseases, among others [5–8]. One example of circadian rhythms in disease can be seen in rheumatoid arthritis (RA), one of the most prevalent autoimmune diseases that affects 0.5% of the global population [9–11]. RA is characterized by chronic destruction of cartilage, leading to severe pain and bone erosion, and is mediated by the over production of proinflammatory cytokines like interleukin-1 (IL-1) and tumor necrosis factor-α (TNF-α), among others [12, 13]. However, inflammation in RA is not continuous but instead fluctuates in circadian patterns, with patients often experiencing a relative increase in peak serum concentrations of multiple cytokines in the early morning (around 2 AM to 7 AM) that corresponds to classically-described “morning stiffness” of the joints [14–17]. Biologic drugs that target proinflammatory cytokines central to RA pathology have been a growing market over the past 25 years [18–20]. However, nearly 20-40% of RA patients fail to respond to many of the available treatments, which are typically delivered at high doses that suppress the immune system and have a number of adverse side effects such as increased risk of infection or cancer [19, 21–23]. Furthermore, such approaches do not account for physiological fluctuations in inflammation [24].

Chronotherapy, also termed “circadian medicine,” represents the delivery of therapeutics to coincide with the body’s circadian rhythm to optimize the efficacy of drug delivery. This emerging approach has demonstrated promising results in conditions like RA [25]. For example, clinical studies have shown that systemic delivery of low dose glucocorticoids with modified release preceding the circadian peak of cytokines improved RA outcomes in comparison to treatment given after the daily peak [26, 27]. Additionally, a pilot study demonstrated that methotrexate, a first-line treatment for RA, taken daily at bedtime improved RA symptoms in comparison to the current practice [28, 29]. In preclinical models, benefit of chronotherapy has been demonstrated for multiple synthetic disease-modifying antirheumatic drugs, including methotrexate, baricitinib, and tacrolimus in collagen-induced arthritis [30–33]. In agreement across both clinical and preclinical studies, aligning medication to the rising phase of daily inflammation led to improved outcomes in comparison to when it was given after peak inflammation had been reached. However, the effect of time of day and application of chronotherapy in biologics remains largely unexplored. Furthermore, circadian drug delivery has several challenges that remain to be addressed, such as daily injections for indefinite periods, or drug administration that is required at inconvenient times of day.

To meet these challenges of long-term circadian drug delivery, we developed synthetic gene circuits to reprogram cells to secrete a therapeutic at a specific time of day by repurposing endogenous circadian mechanisms. Proof-of-concept studies have demonstrated that circadian-driven, or “chronogenetic,” output can be programmed by tapping into the central circadian feedback loop. In the first-generation system, the promoter for *Per2* was cloned upstream of the sequence encoding the biologic interleukin-1 receptor antagonist (IL-1Ra), enabling therapeutic release that fluctuated on a 24-hour period and protected circadian rhythms during IL-1 challenge [34]. Moreover, tissue-engineered cartilage with this chronogenetic circuit was able to align to endogenous circadian rhythms *in vivo* and secrete the biologic on a 24-hour cycle in mice when subcutaneously implanted. Notably, previous research has shown that murine induced pluripotent stem cell (iPSC)-derived cartilage pellets function as well-characterized, reproducible *in vitro* models of articular cartilage inflammation for testing therapeutic gene circuits [35, 36]. As iPSCs are differentiated towards more mature chondrocytes, endogenous circadian rhythms develop and resemble circadian outputs observed in mature primary cartilage [35, 37]. Furthermore, these iPSC-derived cartilage cells can be implanted *in vivo* and function as drug delivery depots that sense and respond to environmental cues [38, 39]. Taken together, the use of gene circuits in tissue engineered cartilage constructs supports a robust approach for investigating and translating tunable chronotherapeutic delivery.

To enable more precisely timed therapeutic prescription, tandem repeats of three different clock-controlled elements (E’-boxes, D-boxes, and RREs, the short DNA sequences that bind circadian transcriptional regulators), were used in this study to drive expression of the therapeutic drug IL-1Ra (**Fig. 1**). In contrast to full-length endogenous promoters, these elements can reduce complexity and increase specificity in expression by excluding non-circadian response elements that control gene expression through other signaling pathways. Importantly, each element naturally targets a unique time for peak expression across a 24-hour period [40]. By developing synthetic gene circuits that drive downstream therapeutic production at different times of the day, “scheduled” drug delivery can be achieved. Ultimately, these systems for drug delivery serve as a platform for translation and assessment of timed biologic delivery that aligns with the endogenous daily fluctuation of cytokine releasing, with the aim of making the therapeutic available when it is most beneficial and reducing the need for exogenous non-physiologic administration of continuous high dose drug treatment.

## Materials and Methods

### Chronogenetic circuit design

Forward and reverse oligonucleotide pairs for each synthetic promoter were acquired from Integrated DNA Technologies (IDT) as custom Ultramer DNA Oligonucleotides. These sequences consisted of the three tandem repeats of *Per2* E’-boxes, *Per3* D-boxes, or *Bmal1* RREs elements [40], followed downstream by a minimal CMV promoter element [36], with SpeI and EcoRI restriction sites on the 5’ and 3’ ends of the oligonucleotide pairs, respectively (Table S1). Forward and reverse pairs were annealed and phosphorylated to form the synthetic promoters. The Per2-GFP-LUC lentiviral plasmid (*Per2* promoter followed by green fluorescent protein, GFP, and luciferase, LUC, separated by a t2a element) was digested with SpeI and EcoRI to remove the existing *Per2* promoter, and then each synthetic promoter was ligated into the backbone to form E’box-GFP-LUC, Dbox-GFP-LUC, or RRE-GFP-LUC [34]. Next, the E’box-GFP-LUC backbone was digested with AgeI and BamHI enzymes to remove GFP. GFP was replaced with the coding sequence for IL-1Ra, which was amplified from the NRE-IL1Ra lentiviral plasmid by PCR to include corresponding restriction sites for AgeI and BamHI (Table S2) [36]. The t2a-LUC region was removed by PCR amplification of the vector excluding this segment, which was ligated to re-circularize the E’box-IL1Ra vector. To form Dbox-IL1Ra and RRE-IL1Ra, the E’box-IL1Ra vector was digested with SpeI and EcoRI to remove the promoter, and the synthetic D-box and RRE promoters with corresponding restriction sites were ligated into the backbone to form Dbox-IL1Ra and RRE-IL1Ra.

### iPSC-derived cartilage differentiation and culture

Murine iPSCs were differentiated to produce cartilage pellets according to a previously developed protocol [41]. First, iPSCs were cultured on mitomycin C-treated mouse embryonic fibroblasts (MEFs) for five days in gelatin-coated dishes. iPSC culture media consisted of Dulbecco’s Modified Eagle Medium-high glucose (DMEM-HG), 20% lot-selected serum, 1% minimum essential medium (MEM) non-essential amino acids, 55 μM 2-Mercaptoethanol, 25 μg/mL gentamicin, and 1,000 units/mL mouse leukemia inhibitory factor (LIF). Following feeder cell subtraction, cells were differentiated towards the mesenchymal linage using high density micromass culture for 14 days in serum-free chondrogenesis media (DMEM-HG; 1% insulin, transferrin and selenous acid+ (ITS+); 1% MEM non-essential amino acids; 1% penicillin/streptomycin; 55 μM 2-Mercaptoethanol; 50 μg/mL ascorbate, and 40 μg/mL proline). On the 3rd and 5th days of micromass culture, media was supplemented with 50 ng/mL bone morphogenic protein 4 (BMP-4) and 100 nM dexamethasone. Micromasses were digested with pronase and collagenase II and plated onto gelatin-coated flasks as pre-differentiated iPSCs (PDiPSCs) and expanded in media with DMEM-HG, 10% lot-selected serum, 1% ITS+, 1% MEM non-essential amino acids, 1% penicillin/streptomycin, 55 μM 2-Mercaptoethanol, 50 μg/mL ascorbate, 40 μg/mL proline, and 4 ng/mL basic fibroblast growth factor (bFGF). Cells were transduced at the PDiPSC stage. Second passage PDiPSCs were harvest and centrifuged in 15-mL conical tubes at a concentration of 250,000 cells per culture. Pellet cultures were maintained for 21 days in media consisting of DMEM-HG, 1% MEM non-essential amino acids, 1% penicillin/streptomycin, 55 μM 2-Mercaptoethanol, 100 nM dexamethasone, 50 μg/mL ascorbate, 40 μg/mL proline, and 10 ng/mL transforming growth factor-β3 (TGF-β3).

For sample collection, cells were synchronized in PDiPSC culture media or pellet culture media, corresponding to their respective differentiation stage, supplemented with 100 nM dexamethasone for one hour, and then cultured in media without dexamethasone or growth factors. Inflammatory challenges were induced by adding 1 ng/mL IL-1β to the culture media after synchronization.

### Lentivirus production and cell transduction

A standard protocol was followed to produce second-generation packaged vesicular stomatitis virus glycoprotein pseudotyped lentivirus [42]. HEK293T cells were transfected with the psPAX2 packaging vector, pMD2.G envelope protein vector, and expression vectors by calcium phosphate precipitation. Media collected from transfected HEK293T cells was filtered and stored at -80°C until use. A viral titer was performed to identify the multiplicity of infection (MOI). For experiments, cells were transduced for 24 hours at the PDiPSC monolayer-stage at an MOI of approximately 3 in media supplemented with 4 μg/mL polybrene.

### Bioluminescence circuit characterization

Following synchronization, pellets were cultured in phenol-free media supplemented with 100 μM luciferin and 0 or 1 ng/mL IL-1β in an enclosed CO_2_-controlled incubator. Luminescence was recorded at 15-minute intervals over 72 hours. Raw data between 12 to 72 hours was normalized and detrended over a 24-hour average with smoothing of 1 point using ChronoStar software [43]. Samples were detrended by dividing the data by the trend to evaluate relative expression change. Period (time required for the signal to repeat), phase (relative shift of the signal), amplitude (height of the signal), and correlation coefficient (CC) to an oscillating curve were recorded as outcomes for circadian assessment. Samples with a period less than 18 hours or greater than 30 hours were excluded from the analysis (n=16 of total samples analyzed); samples with outlying phases were excluded based on Iglewicz and Hoaglin’s robust test for multiple outliers due to poor fit alignment (n=4 of total samples analyzed), and then samples with outlying amplitudes were excluded using the same method (n=8 of total samples analyzed). For sample phases at the cusp of the 24/0-hour timepoint (i.e., D-box samples), 24 hours were subtracted from the phase if the software reported a value greater than 12 hours to allow for linear comparison between groups.

### Gene expression

Gene expression was analyzed using quantitative reverse transcription polymerase chain reaction (RT-qPCR). At each collection timepoint, samples were frozen at -80°C until RNA isolation. To isolate RNA, pellets were homogenized using a miniature bead beater, and then RNA was then isolated according to the manufacture’s protocol (Norgen Biotek). Complementary DNA (cDNA) was produced using Invitrogen SuperScript VILO cDNA master mix. Gene expression was quantified by RT-qPCR using Fast SyBR Green master mix (Applied Biosystems) according to the manufacture’s protocol using the ΔΔCT method to determine relative fold change in gene expression, with respect to endogenous expression of *Gapdh*. Primers for were synthesized by IDT and verified for efficiency (Table S3).

To assess expression kinetics, E’box-IL1Ra, Dbox-IL1Ra, and RRE-IL1Ra pellets were collected at four-hour intervals over 48 hours after synchronization in comparison to respective samples collected at the 0-hour timepoint. Averaged *Il1rn* gene expression at each timepoint was assessed using MetaCycle [44]. Individual samples were sorted by expression at each timepoint and assessed for period by the same methods for sets with values for each timepoint. Overall sinusoidal fit by Prism GraphPad 10 (nonlinear fit for a sine wave with a nonzero baseline, least squares regression, constrained to wavelength <18) was plotted alongside data. To characterize the response to an inflammatory challenge, E’box-IL1Ra, Dbox-IL1Ra, RRE-IL1Ra, and non-transduced pellets were synchronized and cultured with 0 or 1 ng/mL IL-1β. RNA samples were collected 24 and 72-hours post-synchronization and compared with respect to unchallenged non-transduced samples at each timepoint.

### Enzyme-linked immunosorbent assay

Culture media was collected and stored at -20°C until analysis. Concentrations of IL-1Ra in the media were quantified in duplicate using the R&D Systems DuoSet enzyme-linked immunosorbent assay (ELISA) for mouse IL-1Ra with absorbance readings at 450 nm and 540 nm. To assess therapeutic formation over time, individual samples at each timepoint were sorted by concentration to form sample sets, and rate was determined by taking the first derivative of the samples using Prism GraphPad 10 (smoothing with 3 neighbors by 4th order polynomial). Rate was normalized for each sub-column and plotted for data over 6 to 46 hours. Rate of IL-1Ra was assessed using MetaCycle and Prism GraphPad as described for gene expression.

### Biochemical analysis of pellets

Pellets were digested in papain overnight at 65°C. Total sulfated glycosaminoglycan (sGAG) content in the pellets was quantified using a 1,9-dimethylmethylene blue (DMMB) assay and normalized to total DNA content, quantified using a PicoGreen assay (Thermo Fisher) according to the manufacturer’s protocol.

### Histological analysis of pellets

Pellets were fixed in a formalin solution, dehydrated using ethanol, embedded in paraffin, and sectioned into 8-μm thick segments. Pellet cultures were stained with Safranin-O, Fast-Green, and hematoxylin. Brightfield images of slides were taken using an Olympus VS120 microscope at 20x magnification.

### Statistical analysis

Metrics comparing E’-box, D-box, and RRE circadian evaluation were assessed by one-way ANOVA with Tukey’s multiple comparisons test. Circadian metrics for control and therapeutic samples with or without inflammatory challenge were assessed two-way ANOVA with Tukey’s multiple comparisons test. 24- and 72-hour samples with or without inflammatory challenge were evaluated using one-way ANOVA with Sidak’s multiple comparisons test.

## Results

### E’-boxes, D-boxes, and RREs drive circadian output with distinct, prescribed phases

Expression kinetics for each promoter were characterized based on bioluminescent reporter output. In these systems, each synthetic promoter drives the expression of luciferase, which can be continuously monitored to provide real-time expression data. Signals were recorded for 72 hours and reported as normalized, detrended values using a 24-hour moving average, excluding the first 12 hours of data to eliminate artifacts from the addition of the substrate. Therefore, data between 24- to 60-hours post-synchronization was utilized to quantify circadian parameters. Detrended, normalized luminescence from E’-box-, D-box-, and RRE-driven bioluminescent reporters demonstrated circadian periods in the PDiPSC intermediate monolayer stage (E’-box: 26.6±1.1, D-box: 26.4±1.4, RRE: 26.3±1.8 hours) and in the final-stage three-dimensional cartilage pellets that are more representative of articular cartilage (E’-box: 23.9±0.6, D-box: 23.9±1.0, RRE: 23.4±0.5 hours) (**Fig. 2**). As shown in the overlay of circuit output, each promoter drove expression with a unique phase from the fit wave in a 24-hour cycle that was consistent in sequence order for cells at both the PDiPSC and pellet stages. Regardless of phase, there was no significant difference in period between E’-boxes, D-boxes, and RREs for PDiPSCs or pellets. Moreover, these signals showed high correlation coefficients (CC) with the fit that was more consistent between groups with maturation.

**Figure 2.**
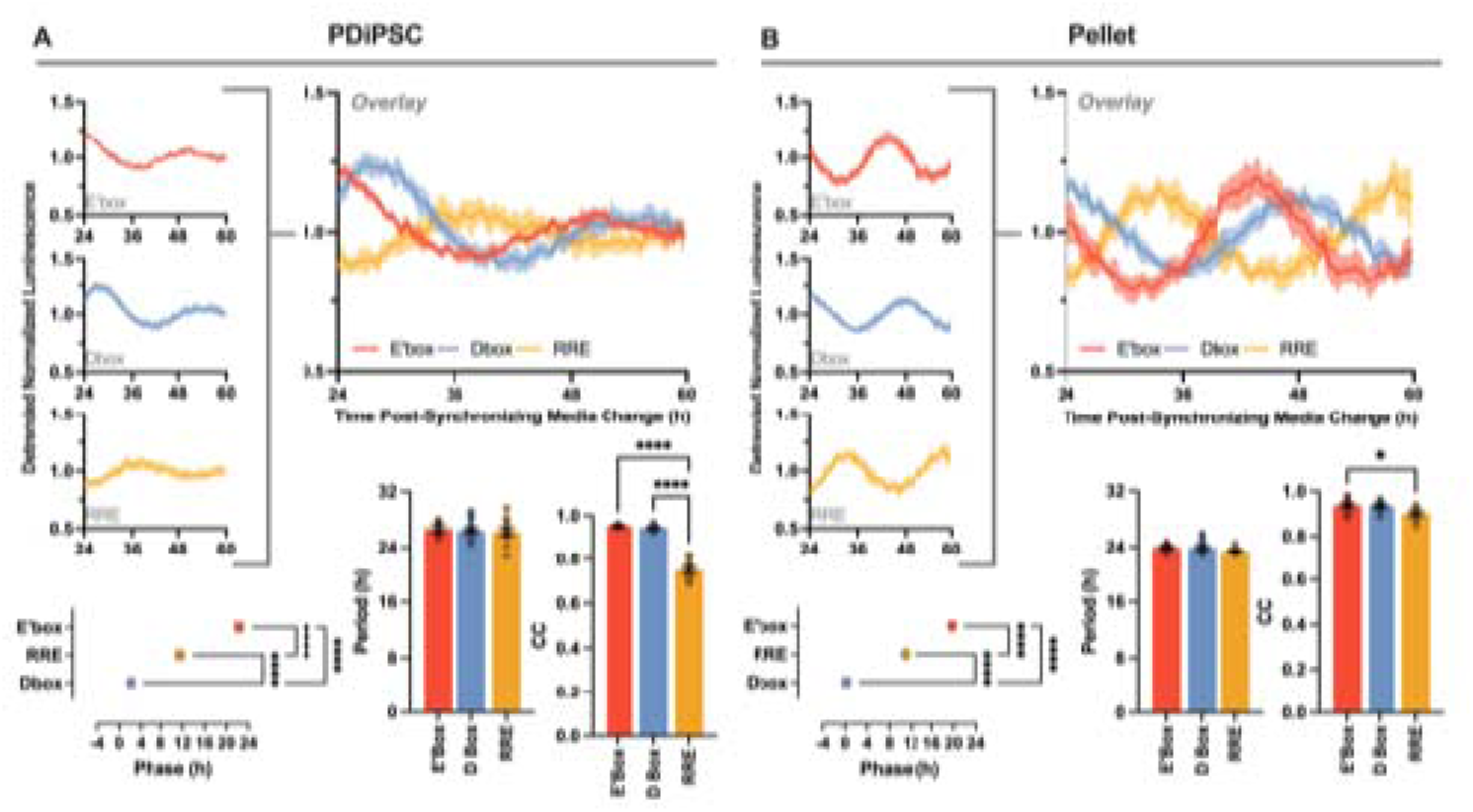
Clock-controlled elements drive circadian output. As promoter elements, E’-boxes, D-boxes, and RREs drive circadian bioluminescence. (A) In PDiPSCs and (B) in cartilage pellets, each promoter drives a reporter that shows a unique phase within a 24-hour period. No significant difference in period is noted between promoters at either stage. With differentiation from PDiPSC to pellet, correlation coefficient (CC) to the sinusoidal fit is more consistent across groups (A, n=5-15/group; B, n=8-10/group; significance by one-way ANOVA with Tukey’s multiple comparisons test, *p<0.05, ****p<0.0001; mean with SEM).

### Chronogenetic circuits drive therapeutic delivery with prescribed phases in a daily cycle

To assess the ability of circuits to functionally generate chronotherapeutic outputs, therapeutic gene and protein expression from E’box-IL1Ra, Dbox-IL1Ra, and RRE-IL1Ra pellets were quantified over 48 hours (**Fig. 3**). Mean *Il1rn* gene expression (Fig, 3A-C) and the rate of IL-1Ra protein formation normalized to the relative production (Fig. 3D-F) for all chronogenetic circuits aligned with sinusoidal curves that had approximately 24-hour periods. Quantification of individual sample sets over a 48-hour period similarly resulted in roughly 24-hour periods for gene expression (E’-box: 22.8±1.3, D-box: 24.4±2.2, RRE: 23.2±0.3 hours) and rate of protein formation (E’-box: 27.6±2.2, D-box: 22.0±1.8, RRE: 24.7±3.4 hours). Based on the curve fit, peak rate of protein formation was estimated to vary in phase in the same sequence as observed for bioluminescence output, peaking at approximately 25 hours for E’-box, 29 hours for D-box, and 31 hours for RRE within the measured time period. Together, this suggests that these circadian promoters support daily drug delivery that can be targeted to peak at distinct phases in a 24-hour period to evaluate IL-1Ra chronotherapy.

**Figure 3.**
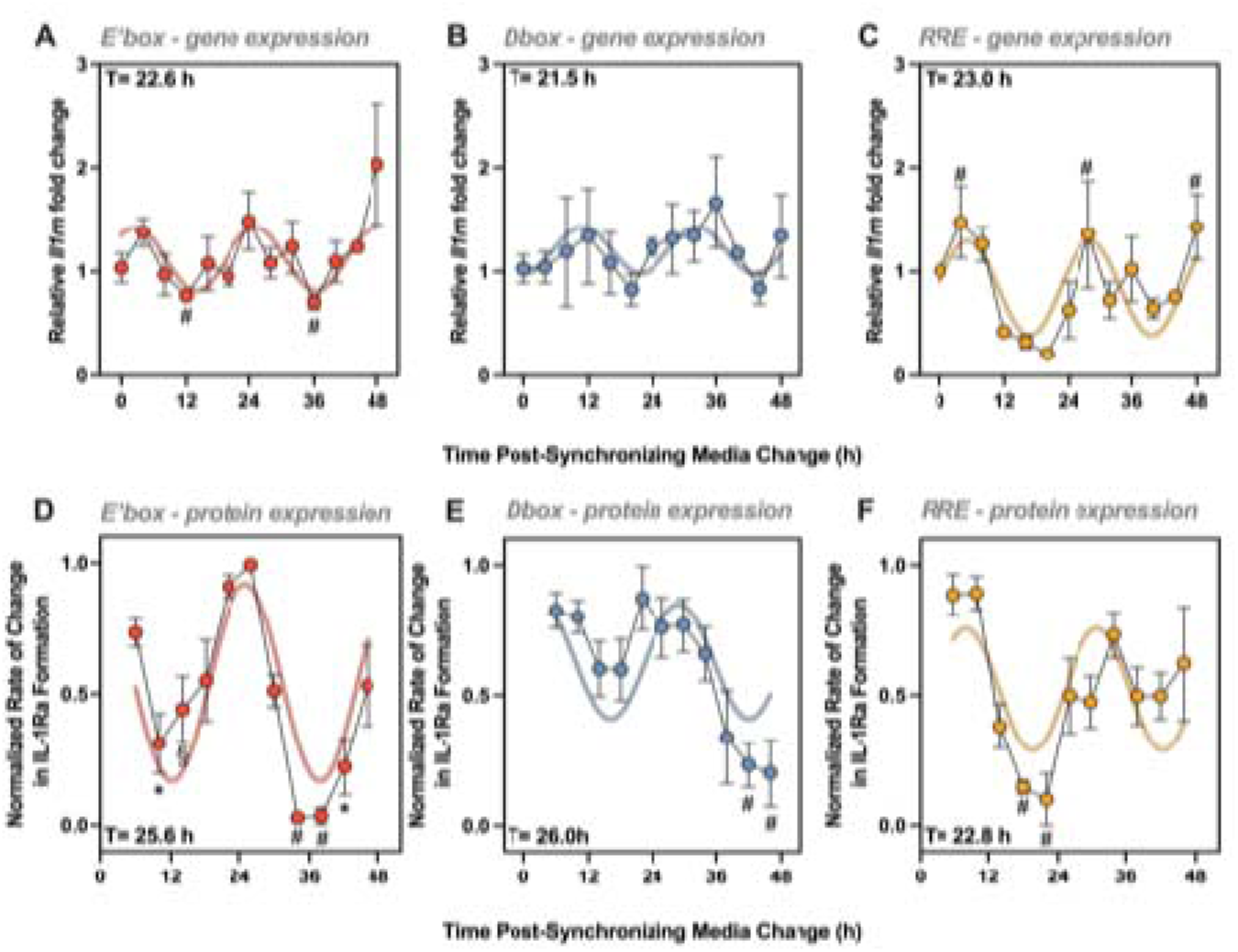
Chronogenetic circuits drive therapeutic delivery. (A-C) Relative fold change in therapeutic *Il1rn* gene expression is cyclical on a daily scale in cartilage pellets (A, ^#^significant from 48h; C, ^#^significant from 20h). (D-F) Rate of therapeutic IL-1Ra protein formation normalized to relative minimum and maximum production by each CCE generates daily-scale drug production. Each promoter drives output with a unique phase in a 24-hour period (D, ^#^significant from 6h, 22h, and 26h, *significant from 22h and 26h, ^significant from 26h; E, ^#^significant from 6h, 10h, and 22h; F, ^#^significant from 6h, 10h, and 24h). Sinusoidal curves with approximate circadian periods (T) fit to both gene expression and rate of protein formation (n=3-6/timepoint; significance assessed by one-way ANOVA with Tukey’s multiple comparisons test, p<0.05 where stated significant; mean with SEM).

### Chronogenetic-driven IL-1Ra protects circadian disruption during an IL-1 inflammatory challenge

To mimic the dysregulated inflammatory conditions in RA, cartilage pellets were subject to a cytokine challenge. Cytokines that are elevated systemically and locally in inflamed RA joints include IL-1β, TNF-α, and IL-6 [45–48]. Since IL-1 is known to disrupts circadian rhythms in cartilage, we evaluated the ability of each therapeutic circuit to protect its respective bioluminescent reporter [35, 49] (**Fig. 4**). Pellets were transduced with only a reporter circuit or with both a reporter and an IL1Ra-generating circuit and evaluated in the presence or absence of an IL-1β challenge to mimic an inflammatory flare. During the inflammatory challenge, E’-box circadian bioluminescence was disrupted, demonstrated by a significant increase in period. However, when E’box-IL1Ra was present, period was not significantly changed between treatment groups. D-box circadian bioluminescence showed a significant decrease in amplitude during challenge while Dbox-IL1Ra mitigated this reduction. Disruption in RRE circadian bioluminescence was characterized by a significant shift in phase, and this shift was reduced with RRE-IL1Ra. Overall, these data support that these therapeutic chronogenetic circuits protect against circadian disruption during inflammatory challenge.

**Figure 4.**
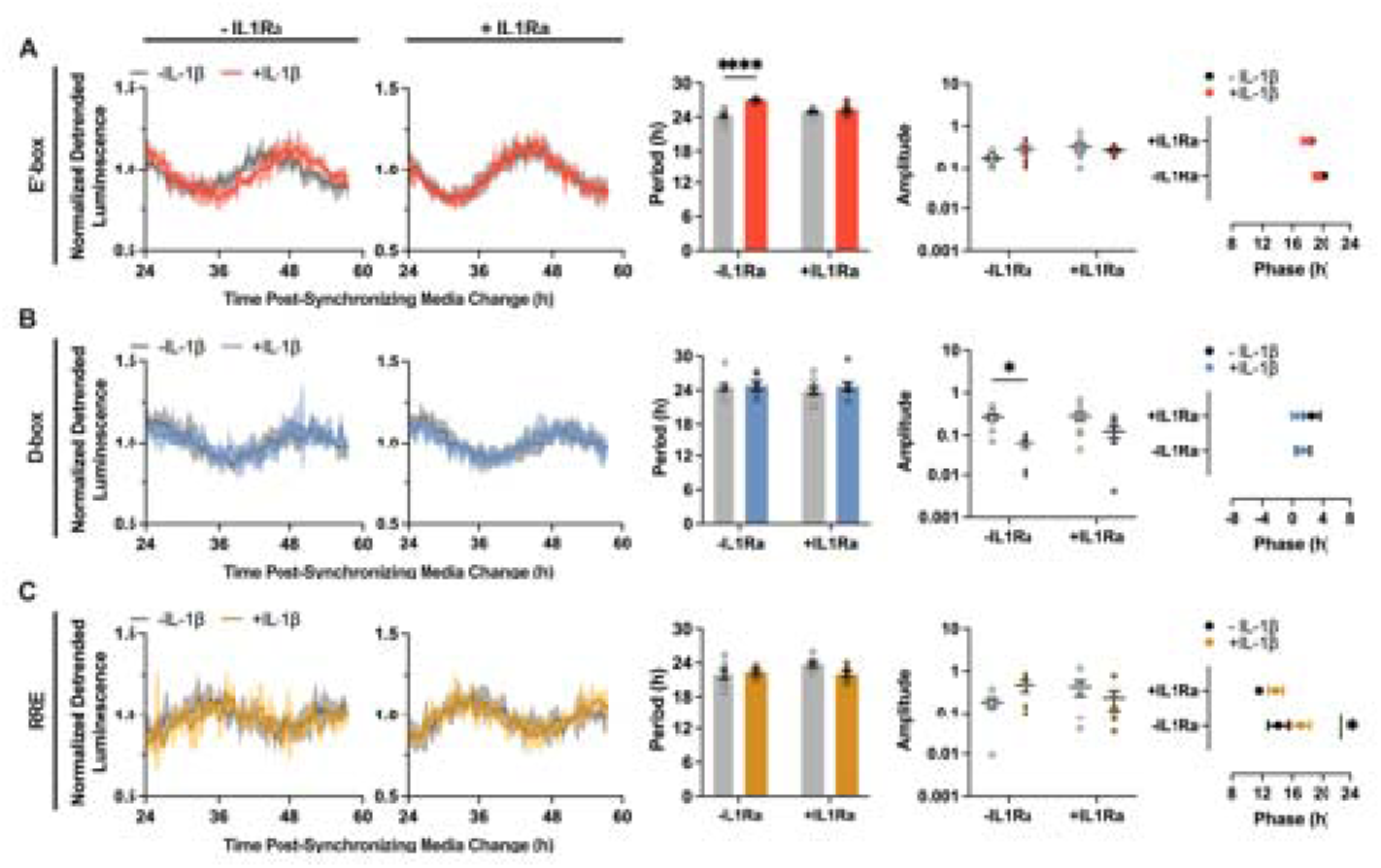
Chronogenetic circuits protect against inflammation-driven circadian disruption. Normalized, detrended luminescence without (left) or with (right) the respective IL-1Ra circuit shows that IL-1β challenge affects the circadian bioluminescence of the reporter but the effect is negated when the therapeutic circuit is present. This is demonstrated by (A) a significantly lengthened period for E’-box, (B) a significantly reduced amplitude for D-box, and (C) a significantly shifted phase for RRE. Regardless of measured circadian disruption, E’box-IL1Ra, Dbox-IL1Ra, and RRE-IL1Ra mitigate these changed to levels that are not significant from unchallenged controls (n=5-9/condition, significance assessed by two-way ANOVA with Tukey’s multiple comparisons test, *p<0.05, **p<0.01; mean with SEM).

### Chronogenetic-driven IL-1Ra protects 3D cartilage pellets as a model of arthritis

To test the therapeutic potential of these chronogenetic circuits in engineered cartilage as an *in vitro* model of arthritis, pellets with E’box-IL1Ra, Dbox-IL1Ra, and RRE-IL1Ra were assessed in the presence or absence of an IL-1β challenge, at shorter and longer timepoints. Between 24- and 72-hours, the concentration of IL-1Ra produced in the culture media increased for all circuits and was not significantly affected by an IL-1β challenge. However, there was a difference between relative concentrations produced by each promoter, with E’-box driving the highest output and RRE driving the lowest output (**Fig. 5A**). Despite this difference in therapeutic concentration, protection of cartilage- and inflammatory-associated gene expression was evident at 24- and 72-hours post-challenge in comparison to non-transduced (NT) control pellets for all circuits (**Fig. 5B**). In non-transduced controls, expression of *Acan* and *Col2a1* decreased approximately 10-fold by 72 hours post-challenge. Pellets with E’box-IL1Ra, Dbox-IL1Ra, and RRE-IL1Ra remained at near baseline levels at both earlier and later timepoints, with no significant difference in expression between challenged and unchallenged groups. Likewise, the expression of *Il6* and *Ccl2* increased approximately 100- to 1000-fold when challenged in non-transduced controls. Inflammatory gene expression for pellets with E’box-IL1Ra, Dbox-IL1Ra, and RRE-IL1Ra was significantly dampened compared to non-transduced groups.

**Figure 5.**
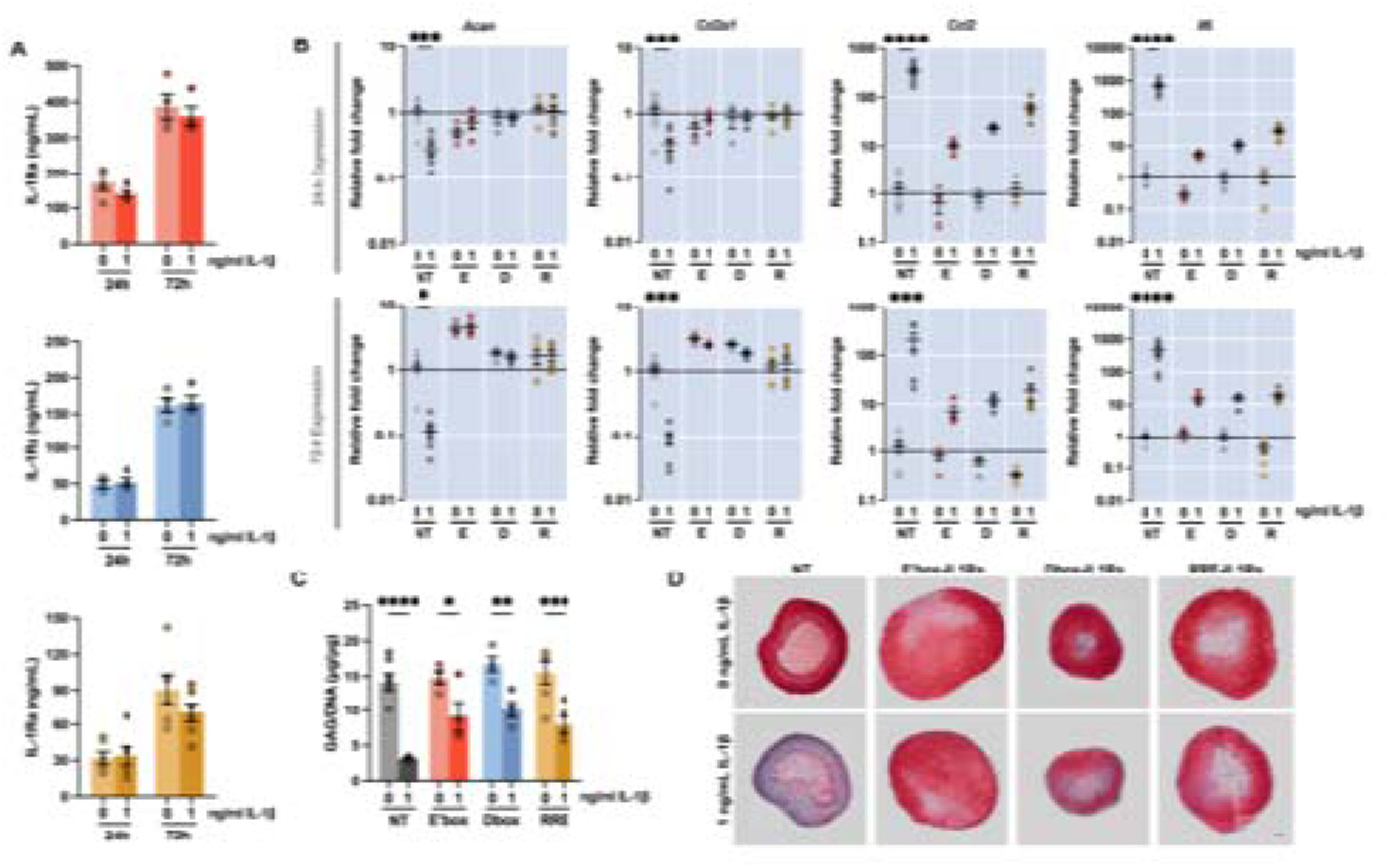
Chronogenetic gene circuits protect against inflammation-driven cartilage damage. (A) IL-1Ra is produced over 72h for all CCE-driven circuits in the presence or absence of an IL-1β inflammatory challenge (n=3-6/condition). (B) For all chronogenetic circuits, IL-1Ra production is sufficient to protect cartilage and dampen inflammatory gene expression during IL-1β challenge in comparison to NT controls at early and late timepoints (n=3-7/condition). (C) Proteoglycan content of cartilage pellets is protected after inflammatory challenge in comparison to NT controls as measured by GAG/DNA content (n=4-7/condition) and (D) representative histological images stained with Safranin-O, Fast-Green, and hematoxylin (n=2-4/condition, scale bar: 100 µm). Significance assessed by one-way ANOVA with Sidak’s multiple comparisons test, *p<0.05, **p<0.01, ***p<0.001, ****p<0.0001; mean with SEM.

Consistent with gene expression data, quantitative measures of sulfated glycosaminoglycans (GAGs) normalized to DNA in pellets showed that non-transduced groups challenged with IL-1β for 72 hours contained approximately one-third the value of this ratio as their unchallenged controls. However, challenged pellets with E’box-IL1Ra, Dbox-IL1Ra, and RRE-IL1Ra retained nearly double the amount of glycosaminoglycans content compared to the challenged non-transduced group, although this was still a significant reduction from their respective unchallenged controls (**Fig. 5C**). Nevertheless, pellets with E’box-IL1Ra, Dbox-IL1Ra, and RRE-IL1Ra displayed a protection in proteoglycan content by histological staining with Safranin-O at 72 hours post-challenge in comparison to non-transduced controls, supporting that these chronogenetic circuits protect the engineered cartilage from inflammation-induced damage at a tissue level (**Fig. 5D**).

## Discussion

“Chronogenetics” is an emerging field that seeks to prescribe controlled gene expression using endogenous cellular timing systems [34]. Here, we applied synthetic chronogenetic gene circuits to develop cell-based constructs that can deliver therapeutic drugs with distinct, prescribed phases within a 24-hour period. Taking a synthetic biology approach, we utilized three unique clock-controlled elements to engineer cell-based strategies for prescribed IL-1Ra production by adapting features of the endogenous circadian feedback loops that maintain cellular timing. We found that three tandem repeats of minimal circadian response elements (E’-boxes, D-boxes, or RREs) adjacent to a basal promoter are sufficient to drive therapeutic IL-1Ra output at three phases within a 24-hour window, increasing our capacity to optimize drug timing with respect to daily inflammatory fluctuations characteristic of RA [14–16, 34]. While we have yet to translate these systems to assess their benefit *in vivo*, they serve as platforms that will support the investigation of timed drug delivery for arthritis and overcome limitations of previous chronotherapy research for RA. Namely, these circuits can target three timeframes that broadly cover a 24-hour period to more precisely evaluate the optimal administration time, act as self-regulated delivery systems that align with natural cycles to eliminate inconvenient or inconsistent delivery times, and extend our study of chronotherapy into biologics, which represent a sizable portion of RA therapeutics [50].

E’-boxes, D-boxes, and RREs are critical regulators of clock gene timing [40, 51]. Tandem repeats of E’-boxes, D-boxes, and RREs are known to oscillate at unique phases within a 24-hour period [40]. Our promoters closely aligned with previous findings, demonstrating that each element targets a unique phase and follows in a sequential order from E’-box to D-box to RRE in a 24-hour period. Characterization of these expression kinetics at the intermediate PDiPSC stage and the mature pellet stage highlights the consistency of this endogenous phase order. Moreover, the increased robustness of the sinusoidal output from PDiPSC to pellet is consistent with previous studies on the development of circadian rhythms throughout maturation of iPSC-derived cartilage [35]. While undifferentiated iPSCs lack rhythmic output of reporters tracking clock gene expression, PDiPSCs demonstrated greater circadian output than iPSCs, and ultimately, mature cartilage pellets had strong oscillations for both *Per2* and *Bmal1* reporters. Our data supports this trend, quantitatively demonstrated by high correlation coefficients to the fit oscillating curve representing the circadian output patterns.

After validating that these minimal promoter elements generated circadian oscillations with unique phases in our iPSC-derived cartilage model, we developed therapeutic chronogenetic circuits to drive IL-1Ra output. As expected, the expression of *Il1rn* downstream of these synthetic promoters resulted in a cyclical relative fold change in gene expression that fit an approximately 24-hour period. Additionally, the rate of IL-1Ra formation over time demonstrated a similar output pattern as gene expression. The phase of the fit curve was consistent in order with bioluminescent output, supporting that the therapeutic generated was driven based on the expected circadian cues.

Next, we sought to understand the impacts of these chronogenetic circuits on circadian rhythms during inflammation-driven disruption. Circadian disruption following IL-1β challenge was characterized primarily by a lengthened period for E’-boxes, a reduced amplitude for D-boxes, and a shifted phase for RREs. IL-1β, a known mediator of RA inflammation, significantly impacted the circadian reporter output observed in cartilage tissue explants and in iPSC-derived cartilage pellets [35, 49, 52]. While the effects of IL-1β challenge on reporters driven by specific clock-controlled elements was uncharacterized, we tested the ability of each therapeutic chronogenetic circuit to protect metrics of circadian disruption for their respective bioluminescence reporters to match the phases of therapeutic output with the phases of bioluminescence. With this experimental design, we noted that the circadian disruptions caused by IL-1β were unique to each promoter.

For these metrics of circadian disruption, the respective therapeutic circuits were largely able to mitigate the observed changes towards baseline levels in our cartilage model. Other work has highlighted that the effect of inflammatory challenge on a circadian disruption depends on the cytokine and its dosage, the cell type assessed, and the clock gene tracked, as well as on the time of challenge relative to the expression of the reporter [49, 53, 54]. In previous reports, *Per2* bioluminescence reporter period in iPSC-derived cartilage pellets was lengthened following IL-1β treatment while amplitude was not significantly affected [35]. The E’-box element, which was derived from the *Per2* promoter, is the main oscillatory driver for *Per2* expression, which aligns with our results showing an increased period for the E’-box-driven reporter during IL-1β challenge [55]. The D-box-driven reporter, which was derived from the *Per3* promoter, displayed a significant decrease in amplitude with IL-1β treatment. Previous studies highlight the importance of D-boxes in maintaining the amplitude of gene expression for both *Per2* and *Per3*, connecting its role to the disruption observed here [55, 56]. Additionally, characterization of clock gene response to cytokines in fibroblasts has demonstrated that IL-1β challenge suppressed *Per3* and *Dbp*, a positive regulator of D-box-mediated transcription [54]. The RRE-driven synthetic promoter was constructed from the RRE element in the *Bmal1* promoter that is considered its key rhythmic driver [57, 58]. In inflammatory conditions that are mediated through the NF-κB pathway, such as IL-6 and TNF-α challenges, changes in the phase of *Bmal1*-driven reporters have been reported, similar to our observation with IL-1β challenge [53, 59]. However, the authors highlight that changes are cytokine- and dose-dependent, suggesting that the specific circadian disruptions vary based on experiment context. Although the mechanisms that underlie how changes manifest as differences in period, amplitude, or phase are not well understood, especially at isolated response elements, physical interactions between inflammatory NF-κB and clock proteins have been suggested to affect binding availability that could lead to the observed changed in circadian metrics [53, 60]. Regardless of the metric assessed, measurements for period, amplitude, and phase disrupted in E’-box, D-box, and RRE-driven circuits, respectively, were all improved to levels similar to controls with therapeutic circuits, highlighting the therapeutic potential of this delivery approach.

At a tissue-level, we further tested the capacity of each circuit to protect cartilage pellets from IL-1-induced degradation. This model system permits rapid assessment of inflammatory- and cartilage-associated shifts in gene and protein expression following a challenge, which are evident over 72 hours [36]. First, we characterized the concentration of IL-1Ra that each circuit was secreting into the culture media at 24- and 72-hours post-challenge and noted that E’-box-, D-box-, and RRE-driven outputs were not equivalent. This was not unexpected, as we have previously shown that the level of *Per2* and *Bmal1*-driven reporter output differed in mature cartilage pellets [35]. Average *Per2*-driven detrended bioluminescence was nearly 10-fold higher than average *Bmal1*-driven output, aligning with the present findings that E’box-IL1Ra (derived from *Per2* promoter) generated greater therapeutic output than RRE-IL1Ra (derived from *Bmal1* promoter) [35, 40]. Therefore, we hypothesize that the difference in therapeutic output from each promoter is due to its endogenous regulation. In summary, our analysis highlights the capacity of these systems to generate sufficient therapeutic to have a measurable impact on an *in vitro* model of RA.

Despite the differing levels of therapeutic output generated by these circuits, the concentration of secreted IL-1Ra was not significantly impacted by IL-1β challenge and was sufficient to dampen inflammatory damage in our *in vitro* model. While this can be benchmarked using constitutively active IL-1Ra or media dosed with recombinant IL-1Ra, we sought to compare the effect of chronogenetic circuits to their non-transduced controls as a representation of native cartilage in the RA environment [36, 61]. Moreover, as a translational approach, constant systemic delivery of IL-1Ra is undesirable since it can worsen arthritis outcomes, supporting the need for tailored delivery systems [62]. For effective inhibition of the inflammatory effects of IL-1, it is estimated that 10- to 100-fold excess of IL-1Ra over IL-1 is necessary [63–65]. At the 0-hour timepoint, an IL-1β challenge was initiated at 1 ng/mL in fresh culture media. By 24 hours, the concentrations of IL-1Ra in the culture media were approximately 30, 50, and 150 ng/mL for RRE-IL1Ra, Dbox-IL1Ra, and E’box-IL1Ra, respectively, representing fold changes over IL-1β likely to dampen the inflammatory response. Moreover, by 72 hours post-challenge, this concentration had risen to more than double this amount. Protection was evident by increased *Acan* and *Col2a1* expression, genes typical of healthy articular cartilage, and dampened *Ccl2* and *Il6* inflammatory gene activation in comparison to non-transduced controls. Likewise, protection in proteoglycan content was quantitatively and qualitatively shown by a reduced loss of sulfonated glycosaminoglycans normalized to DNA content and a strong red staining with Safranin-O following IL-1β challenge. Together, these data support chronogenetic circuits as a promising therapeutic strategy that produce the biologic IL-1Ra to mitigate cartilage degradation in an *in vitro* model of RA.

As a therapeutic option, recombinant IL-1Ra, or anakinra, is clinically approved as a biologic disease-modifying antirheumatic drug but has shown low efficacy and is limited by its requirement for a relatively high-dose daily injection due to its short half-life [66] and generally uncontrolled timing of delivery. In clinical trials, anakinra was considered modestly efficacious for RA treatment, with the most common adverse side effect being injection site reactions [67]. To overcome the burden and adverse effects of daily injection, we have developed a cell-based drug delivery system that can endogenously generate therapeutic output based on internal circadian cues for specifically timed therapeutic delivery that uses the short half-life of IL-1Ra as an advantage for daily-scale chronotherapy. Previous work has shown that iPSC-derived cartilage that is genetically edited to respond to inflammatory cues for “smart” therapeutic release can be implanted peripherally *in vivo* and produce significant amounts of IL-1Ra for therapeutic impact [38, 39]. Similarly, these chronogenetic designs can be translated into an implantable or injectable cell-based delivery system that provides daily doses of a therapeutic without repeated injections. Importantly, we have previously demonstrated that circadian-driven drug delivery can align to an animal’s circadian rhythm for multiple weeks and automatically adjust daily delivery when shifted to a reversed light-dark cycle [34]. The circuits developed in this study support increased flexibility to target multiple phases in a 24-hour window, allowing future studies to tune delivery to best dampen daily cytokine fluctuations for optimal arthritis treatment. While this strategy holds promise for bridging the gap of translating biologics into chronotherapy, this hypothesis remains to be tested, representing a critical next step for this research. *In vitro* analysis demonstrates proof-of-concept support for phase-specific chronogenetic gene circuits but cannot recapitulate the complex, multi-cytokine circadian fluctuations in inflammation *in vivo*. While implantation of cartilaginous constructs has demonstrated benefit for the translation of other synthetic gene circuits, challenges that could arise include differences in the expression profiles of circadian transcriptional response elements between our observed *in vitro* results and the *in vivo* output, inability to generate adequate dosing, or complications with long-term immune compatibility of implantable constructs. Additionally, the application of this strategy to more commonly prescribed biologics, such as TNF or IL-6 inhibitors, remains to be investigated.

Beyond RA, numerous conditions fluctuate in severity across the daily cycle, including those that affect the cardiovascular, gastrointestinal, and musculoskeletal systems, among many others [68]. The timing of daily medication to when it is most effective has already shown clinically relevance in the treatment of conditions ranging from hypertension to cancer [69, 70]. This phenomenon is not surprising, as both disease mediators like immune activity and metrics that determine drug pharmacokinetics (e.g., metabolism) are dependent on time of day [71, 72]. However, feasibility and compliance are likely low if daily administration is most efficacious only when given at an inconvenient time, e.g., the peak in RA inflammatory mediator release occurs around 2 AM necessitating patients to rise in the middle of the night to take their medication [26, 27]. Therefore, a living implant that can time medication prescription for the optimal circadian phase may be able to alleviate the need for exogenous daily injection and potentially improve medication compliance as well as treatment outcomes. By using clock-controlled elements that target unique phases in the 24-hour cycle, our system can be personalized to an individual’s circadian rhythm for optimal self-regulated chronotherapy.

## Data, Materials, and Software Availability

All study data are included in the article. No new software was developed for this study.

## Funding

This work was supported by the Shriners Hospitals for Children and the National Institutes of Health (AG15768, AG46927, AR080902, AR072999, AR073752, AR074992, AR078949).

## Author Contributions

AC, LP, CP, EH, and FG developed the concept of the study; AC, CP, EH, and FG designed the studies; AC, FP, and OO performed the studies and analyzed the data; AC and FG wrote the manuscript; all authors edited and approved the manuscript.

## Competing Interest Statement

FG is an employee and shareholder in Cytex Therapeutics, Inc. FG and LP have filed intellectual property on topics related to the content of this study (US Patent App. 18/284,487, 2024). The authors declare no other competing interests.

